# A Reevaluation of the Effect of Dietary Restriction on Different Recombinant Inbred (RI) Lines of Male and Female Mice

**DOI:** 10.1101/2021.06.25.449984

**Authors:** Archana Unnikrishnan, Stephanie Matyi, Karla Garrett, Michelle Ranjo-Bishop, David B Allison, Keisuke Ejima, Xiwei Chen, Stephanie Dickinson, Arlan Richardson

## Abstract

Dietary restriction (DR) was reported to either have no effect or reduced the lifespan of the majority of the 41-recombinant inbred (RI)-lines studied (Liao et al., 2010). In an appropriately power longevity study (n > 30 mice/group), we measured the lifespan of the four RI-lines (115-RI, 97-RI, 98-RI, and 107-RI) that were reported to have the greatest decrease in lifespan when fed 40% DR. DR increased the median lifespan of female and male 115-RI mice and female 97-RI and 107-RI mice. DR had little effect (less than 4%) on the median lifespan of female and male 98-RI mice and male 97-RI mice and reduced the lifespan of male 107-RI mice over 20%. While our study was unable to replicate the effect of DR on the lifespan of the RI-mice (except male 107-RI mice) reported by Liao et al. (2010), we found that the genotype of a mouse had a major impact on the effect of DR on lifespan, with the effect of DR ranging from a 50% increase to a 22% decrease. No correlation was observed between the changes in either body composition or glucose tolerance induced by DR and the changes observed in lifespan of the four RI-lines of male and female mice. These four RI-lines of mice give the research community a unique resource where investigators for the first time can study the anti-aging mechanism of DR by comparing mice in which DR increases lifespan to mice where DR has either no effect or reduces lifespan.

**Graphical Abstract:** 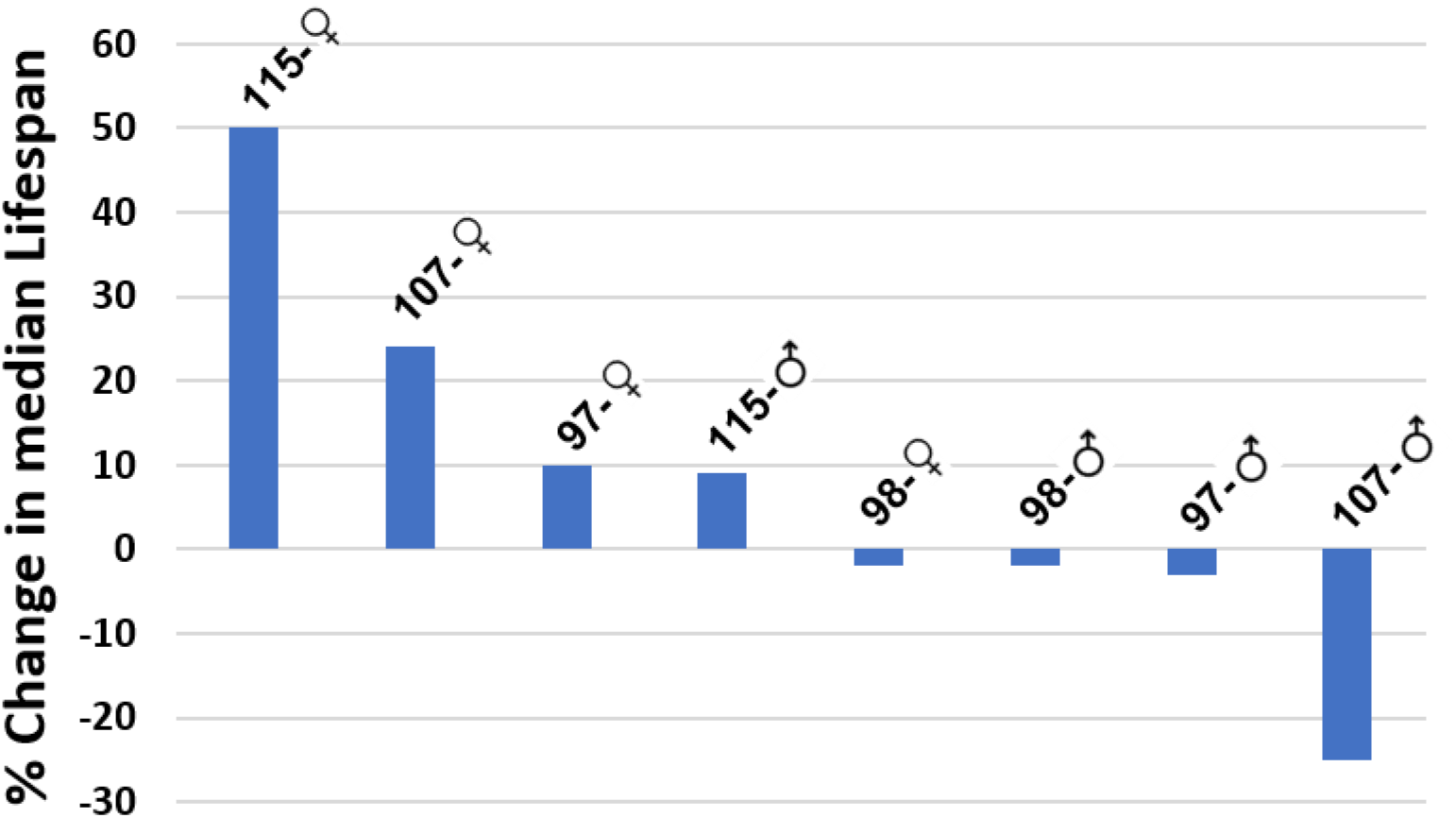

**Brief Abstract:** In this study we reevaluate the effect of the genotype and sex of the recombinant inbred strains of mice on their response to dietary restriction. Out of the eight lines studied, half of the groups showed an increase in lifespan when the diet was restricted by 40% and the other half either did not respond or showed a decrease in lifespan. The recombinant inbred lines used in this study are potentially important resource for aging research to identify pathways and understand the mechanism involved in the anti-aging action of DR.

## Introduction

The first and the most studied manipulation shown to increase lifespan in mammals is dietary restriction (DR). The classic study by McCay et al. in 1935 showed that one could increase the lifespan of rats by dramatically reducing their food consumption early in life. Since this initial observation, numerous laboratories have confirmed these results and have shown that reducing food consumption 30 to 50% (without malnutrition) consistently increased both the mean and maximum lifespans of both laboratory rats and mice (Weindruch and Walford, 1988; Masoro, 2005). The increase in lifespan by DR is similar for laboratory rats and mice used in aging research and similar for females and males, i.e., no pronounced sexual dimorphism was observed (Turturro et al., 1999; Austad, 2017), which is different than has been reported for other manipulations that the Intervention Testing Center has shown to increase the lifespan of mice. The exception is DBA2 mice where DR increased the lifespan of female mice twice as much as male mice.

The effect of DR on longevity is not limited to rodents as DR has been reported to increase the lifespan of a large number of diverse animal models in addition to rodents: invertebrates (Kapahi et al., 2017), dogs (Lawler et al., 2005), and non-human primates (Mattison et al., 2017; Pifferi et al., 2018). Because of the broad effect of DR on lifespan, it has become accepted that the effect of DR on lifespan is universal, i.e., it occurs in all organisms. However, the universality of DR’s effect on longevity was called into question in 2010 when Liao et al. reported the effect of 40% DR on the lifespans of 41 different recombinant inbred (RI) lines of female and male mice. Surprisingly, less than one-third of the RI-lines showed a significant increase in lifespan as was expected. On the other hand, approximately one-third of the RI-lines mice showed a decrease in lifespan on the DR diet and one-third showed no effect of DR on lifespan. These data were a surprise to the research community because they contradicted the prevailing view that DR was a universal, beneficial intervention with respect to lifespan and aging. However, a few previous studies, which had largely gone ignored, also reported that some mouse strains did not show an increase in lifespan when fed a DR diet, e.g., male wild-caught mice (Harper et al., 2006) and male DBA/2 mice (Forster et al., 2003), although Turturro et al. (1999) showed DR increased life span of male DBA/2 mice. In addition, Mattison et al. (2012) reported that DR did not significantly increase the lifespan of rhesus monkeys.

One of the major limitations of the study by Liao et al. (2010) was the number of mice used to measure lifespan in each RI-line of male and female mice, which was limited to less than 10 mice per group and in many cases only 5 mice. Therefore, the goal of this study was to determine the replicability of the lifespan data for the RI-lines of mice when a larger number of mice (e.g., 30 to 45 mice/group) was used to assess the effect of DR on lifespan. Because we could only study a limited number of strains of mice using larger numbers of mice to measure lifespan, we focused our attention on those RI-lines that were reported to show a decrease in lifespan for the following reasons. First, we felt that the data from the RI-lines that showed no significant increase in lifespan could have simply resulted from the small number of mice studied, resulting in the inability to detect a significant difference in lifespan. Therefore, we felt it was more likely that the RI-lines showing a decrease in lifespan would give us the best opportunity to identify RI-lines that did not respond to DR. Second, we were interested in determining if DR actually resulted in a decrease in the lifespan of the RI-lines because such an observation is rare, and in many cases where it has been observed, it has not been replicated. We describe below the effect of DR on the lifespans of male and female mice from four RI-lines of mice: 115-RI, 107-RI, 98-RI, and 97-RI. Our data show that four out of the eight groups of mice studied showed a significant increase in lifespan with DR while the other four show either no significant effect of DR on lifespan or reduced lifespan.

## Results

### Lifespan Analysis

One possible explanation for the contradictory data on the effect of DR on the lifespan of the RI-mice could arise because the level of DR required to increase lifespan is genotype-dependent. In other words, it is possible that 40% restriction used by Liao et al. (2010) had a negative effect on lifespan of some of the RI lines. Therefore, a lower level of DR might increase the lifespan of the genotypes that did not respond or responded negatively to DR. This possibility is supported by two studies that showed lower levels of DR are effective at increasing the lifespan of rats (Richardson et al, 2016) and mice (Mitchell et al., 2016). To test this possibility, we first studied the effect of various levels of DR (10, 20 and 40%) on the RI-line that Liao et al. (2010) reported DR to have the greatest negative effect on lifespan, 115-RI mice, e.g., DR (40%) reduced the mean survival of female and male 115-RI mice ~85% and ~70%, respectively. Figure 1 shows the lifespan curves we obtained from the female and male 115-RI mice fed *ad libitum* (AL) or the three levels of DR, and Table 1 gives the lifespan data and the statistical analysis of these data. It is apparent that the lifespan of the female 115-RI mice is much shorter than the male mice, e.g., median lifespan is ~30% less for female mice compared to male mice. Liao et al. (2010) also reported a similar difference in the lifespan of male and female 115-RI mice. As can be seen from Figure 1A and Table 1, 40% DR significantly increased the lifespan of both female and male mice whether measured by the mixed effects Cox models or the parametric models with Gompertz distribution. However, DR had a much greater effect on the lifespan of the female 115-RI mice than male mice, e.g., median survival was increased 50% for female mice compared to only 9% for males. DR (40%) also significantly increased both the median and mean survival of the female 115-RI mice; however, the increase in the median or mean survival of the male 115-RI mice was not statistically significant. The 90^th^ percentile survival was increased by 35% and 12% for female and male mice, respectively, fed 40% DR. We also used the maximum lifespan test developed by Gao et al. (2008) to statistically test for significant differences in maximum lifespan by testing for differences in the upper tail of the distribution in the survival data. The maximum lifespan test did not quite reach statistical significance for the female 115-RI mice but was significant for the male 112-RI mice.

**Figure 1.**
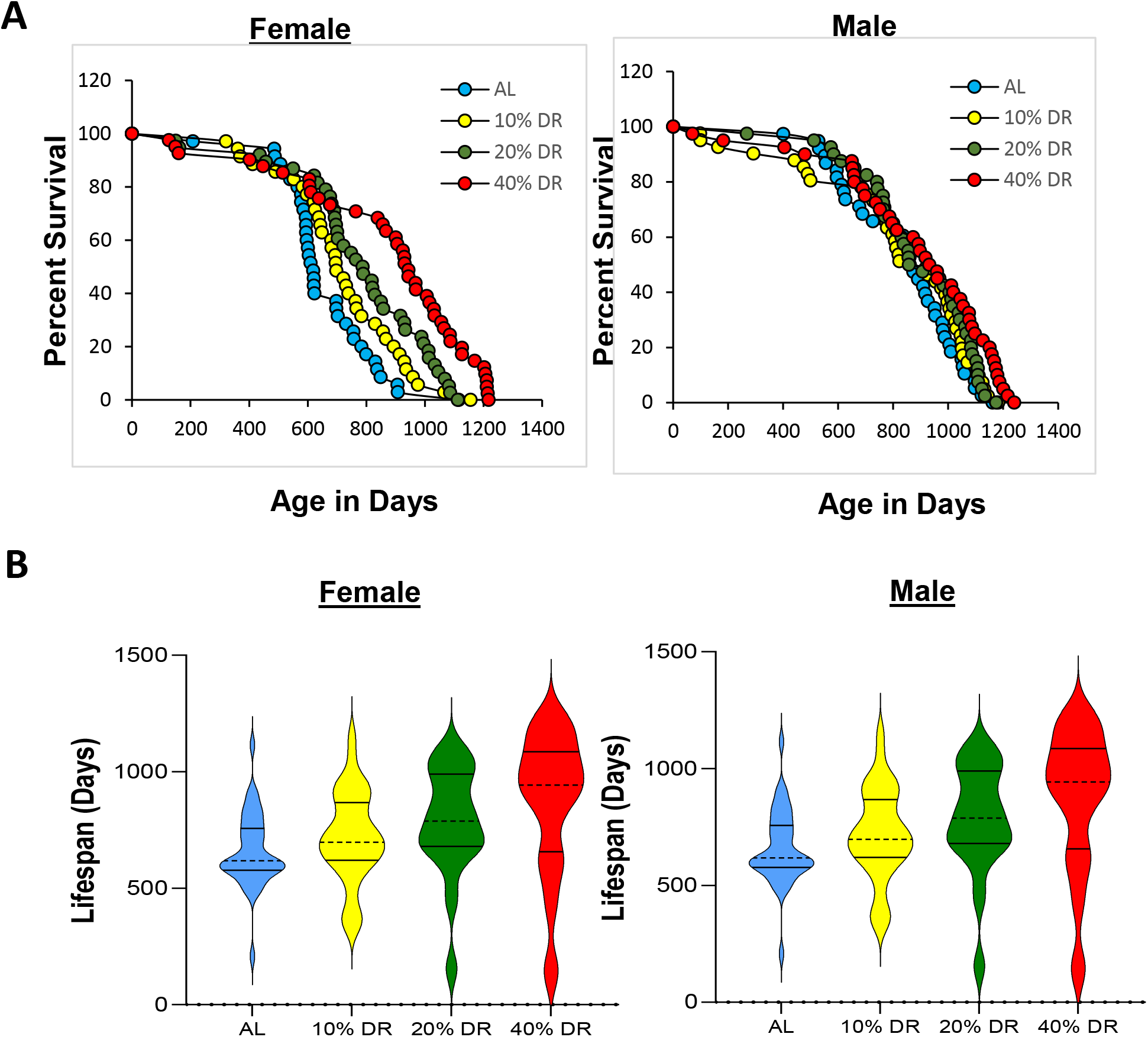
Lifespan of female and male 115-RI mice fed AL and DR. **Panel A** shows the Kaplan-Mier survival curves for mice fed AL (blue) and 10 (yellow), 20 (green), and 40% (red) DR. The number of mice in each group and the analysis of the survival data are given in Table 1. **Panel B** shows the violin plots for the distribution of the lifespans for the age at death for each of the mice in the four groups of 115-RI mice. The solid lines show the quartiles and the dashed line the median.

**Table 1.**
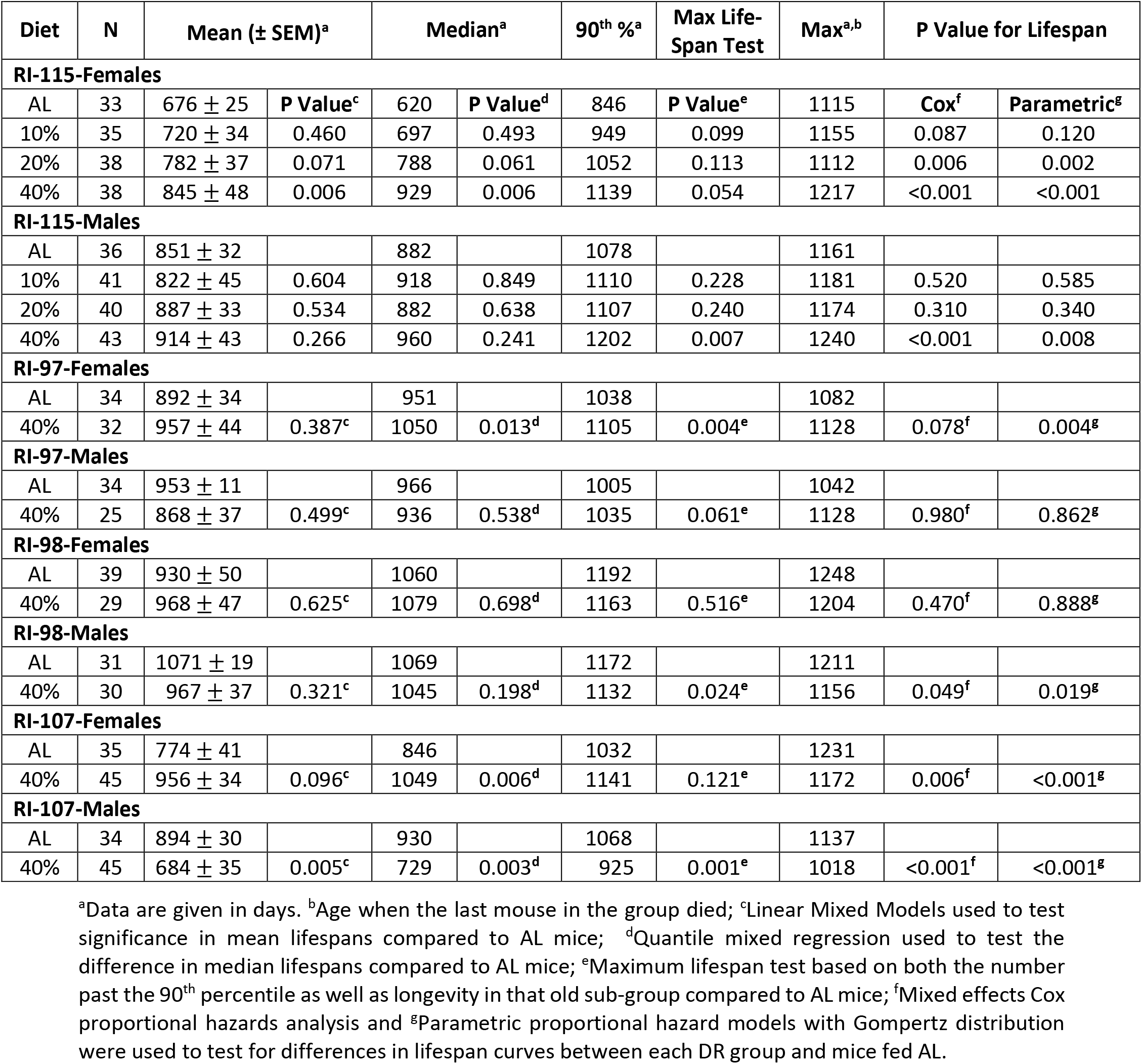
Effect of DR on the Lifespan of Female and Male 115-RI Mice.

As can be seen from Figure 1 and Table 1, lower levels of DR had a smaller effect on lifespan than 40% DR such that in male 115-RI mice, 10 and 20% DR did not significantly increase any measure of lifespan. In contrast, the survival curves for 10, 20, and 40% DR show a graded effect of DR on lifespan of female 115-RI mice, i.e., greater the level of DR the greater the increase in survival. A similar trend was observed when the lifespan data were presented as violin plots (Figure 1B). The lifespan of the female 115-RI mice was significantly increased by 20% DR as measured by either the mixed effects Cox models or the parametric models, resulting in a 16%, 27%, and in 24% increase in mean, median, and median and 90^th^ percentile survival, respectively. However, the increase in mean and median survival was not significant and the maximum lifespan test was not significant. Although 10% DR increased both the median and 90^th^ percentile survival of the female 115-RI mice by 12%, none of the measures of lifespan were significantly increased by 10% DR.

Because 10 and 20% DR did not show any evidence of a greater increase in lifespan of the 115-RI mice compared to 40% DR, we focused our effort on the effect of only 40% DR on the lifespan, which allowed us to study the effect of DR on three other RI lines: 97-RI, 98-RI, and 107-RI mice. Liao et al. (2010) reported that the mean survival of both female and male 97-RI mice were reduced over 50% by 40% DR. As Figure 2A and Table 1 show, we found that 40% DR significantly increased the lifespan of female 97-RI mice as measured by the parametric hazard analysis, as well as a significant (10%) increase in median survival. DR increased the 90^th^ percentile survival 6%, and the maximum lifespan test was also significant. The violin plots in Figure 3A also show a shift toward the DR mice living longer. In contrast, 40% DR had no significant effect on any measure of the lifespan of male R97-RI mice. However, as can be readily observed from the survival curves in Figure 2A or the violin plots in Figure 1S, DR resulted in an increase in deaths in the first half of life in the male 97-RI mice; however, the survival was similar in the later-half of the lifespan.

**Figure 2.**
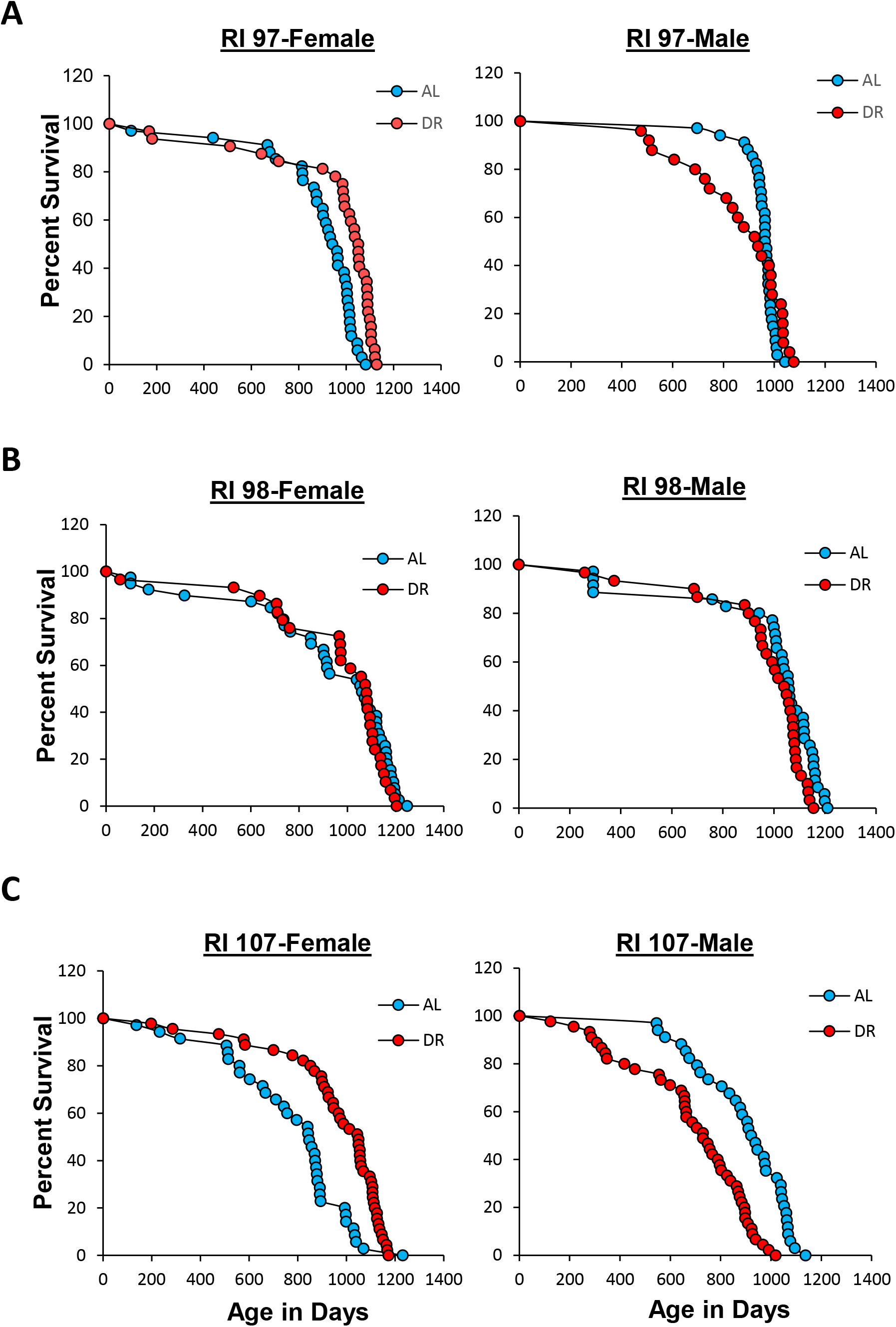
Effect of 40% DR on the lifespan of female and male RI mice. **Panel A, Panel B, and Panel C** shows the Kaplan-Mier survival curves for female and male 97-RI, 98-RI, and 107-RI mice, respectively, fed AL (blue) and 40% (red) DR. The number of mice in each group and the analysis of the survival data are given in Table 1.

**Figure 3.**
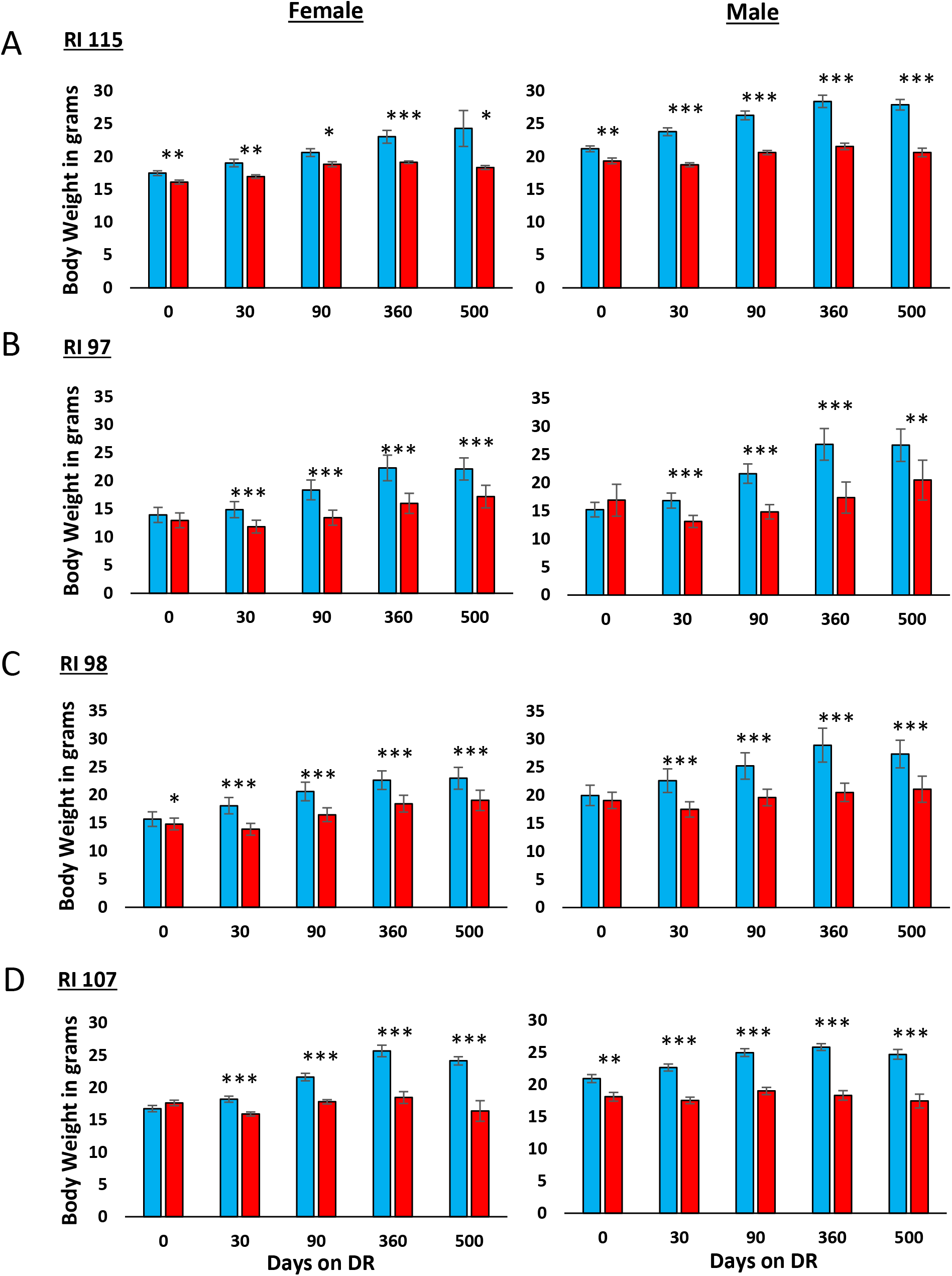
Effect of 40% DR on the body mass of female and male RI mice. The data show the mean ± SEM for 8-10 animals per group except for 97 Male DR (time points 360 and 500) and 107 male and Female DR (time point 500) groups which have 5 mice. **Panel A** 115-RI mice**, Panel B** 97-RI mice, **Panel C**, 98-RI mice, and **Panel D** 107-RI mice. The data for each time point was analyzed as AL Vs DR by twotailed students t-test. Values where the DR mice (red bars) are significantly different from AL mice (blue bars) are shown by *p>0.05, **p.0.01, and ***p>0.001.

We next studied 98-RI mice because Liao et al. (2010) reported that 40% DR reduced the mean survival of both female and male 98-RI mice over 40%. The survival curves and lifespan data for the female and male 98-RI mice in Figure 2B and Table 1 show that 40% DR had no significant effect on the lifespan of the female mice. On the other hand, we observed a statistically significant decrease in the lifespan of male 98-RI mice as measured by both the mixed effects Cox models and the parametric models. The decrease in mean, median lifespan, and 90^th^ percentile of 10%, 2%, and 4% was small and not significant for mean and median lifespan. However, the maximum lifespan test showed a significant difference for the DR mice compared to AL mice. The violin plots in Figure 1S in the supplement also show that the distribution of the lifespan data is similar for AL and DR in both female and male 98-RI mice.

The 107-RI mice were the last RI line we studied. We selected these mice because Liao et al. (2010) reported that this RI-line showed one of the greatest sex differences in the effect of 40% DR on lifespan. DR had no effect on the lifespan of female 107-RI mice but reduced the mean survival of male 107-RI mice by 50%. The lifespan data in Figure 2C and Table 1 show that 40% DR increased the lifespan of female 107-RI mice as measured by either the mixed effects Cox models or the parametric models, resulting in a 24% increase in both mean and median survival and a 10% increase in 90^th^ percentile survival. The increase in median survival was significant; however, either the change in mean survival or the maximum lifespan test were statistically significant. In contrast, 40% DR resulted in a statistically significant decrease in the lifespan of male 107-RI mice as measured by either the mixed effects Cox models or the parametric models, resulting in a 22-23% decrease in the mean and median survival, which was statistically significant for both. A 13% decrease in the 90^th^ percentile survival was observed, and there was a statistically significant difference as measured by the maximum lifespan test. The violin plots in Figure 1S also show that DR shifted the distribution of lifespan of female 107-RI mice to longer lifespan while DR shifted the distribution to a shorter lifespan in male mice.

### Analysis of body mass/composition and glucose tolerance

In their study with the RI mice, Liao et al. (2011) reported that the effect of 40% DR on lifespan was inversely correlated fat reduction, i.e., mice showing the lowest reduction in fat when fed 40% DR were more likely to have extended lifespan. In comparing male and female C57BL/6 and DBA/2 mice fed 20% and 40% DR, Mitchell et al. (2016) also found that the mice that preserved their fat mass in response to DR showed the greatest increase in survival. Therefore, we measured the effect of DR on body and fat mass in the four RI lines of male and female mice to determine if changes in body mass or composition were correlated with the ability of DR to increase the lifespan of the mice. The data in Figure 3 shows the body weights of the four RI-lines of mice fed AL or 40% AL. As expected, all of the mice showed a decrease in body weight. When measuring body composition, we observed no significant change in the percent of lean body mass with DR in most of the RI-lines at any age (Figure 2S in supplement). However, the changes in fat composition by DR varied in the four RI-lines. Figure 3S in the supplement shows the fat mass in grams and Figure 4 shows the percent fat for the four lines from ~2 to 18 months of age. The percent body fat was reduced in male and female 115-RI mice and in male 97-RI and 107-RI mice. On the other hand, DR did not reduce the percent body fat in the female 97-RI and 107-RI mice and male 98-RI mice. Interestingly, the percent body fat was significantly increased in the female 98-RI mice. To determine if there was correlation between the change in body mass or composition induced by DR and lifespan, we plotted the percent change in fat mass, body mass, and lean body mass induced by DR at 12 and 18 months of age versus the change in medium lifespan induced by DR. As shown in Figure 5, we found no significant correlation between the changes in fat mass and lifespan. Interestingly, the group (female 98-RI mice) that showed the least change (actually slight increase) in fat mass by DR showed no increase in lifespan by DR. We also observed no correlation between changes in body mass or lean body mass and lifespan.

**Figure 4.**
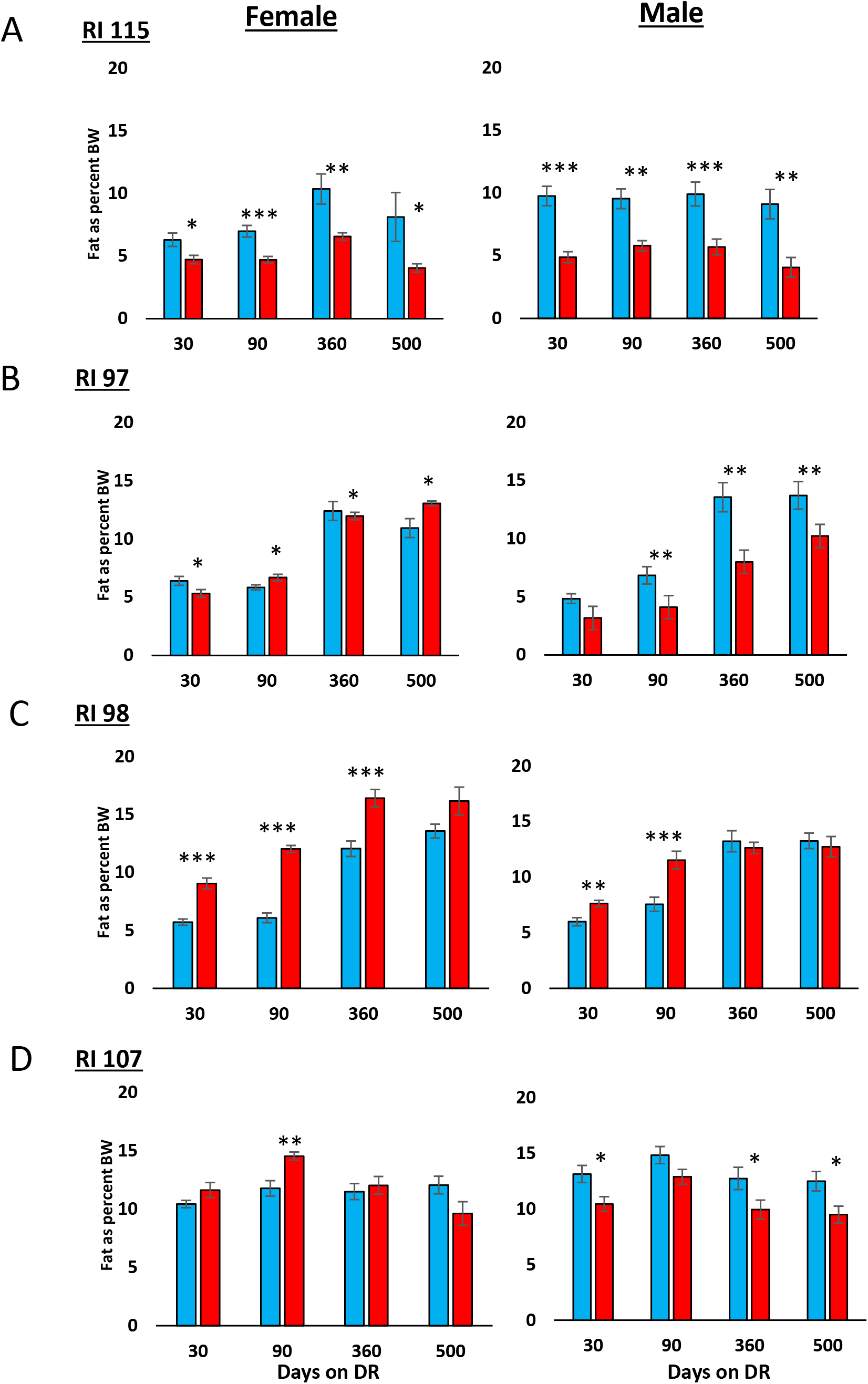
Effect of 40% DR on the percent fat mass of female and male RI mice. The percent fat mass is expressed as the fat mass divided by the body weight of each animal, and the data show the mean ± SEM for 8-10 animals per group except for 97 Male DR (time points 360 and 500) and 107 male and Female DR (time point 500) groups which have 5 mice. **Panel A** 115-RI mice**, Panel B** 97-RI mice, **Panel C**, 98-RI mice, and **Panel D** 107-RI mice. The data for each time point was analyzed as AL Vs DR by two-tailed students t-test. Values where the DR mice(red bars) are significantly different from AL mice (blue bars) are shown by *p>0.05, **p.0.01, and ***p>0.001.

**Figure 5.**
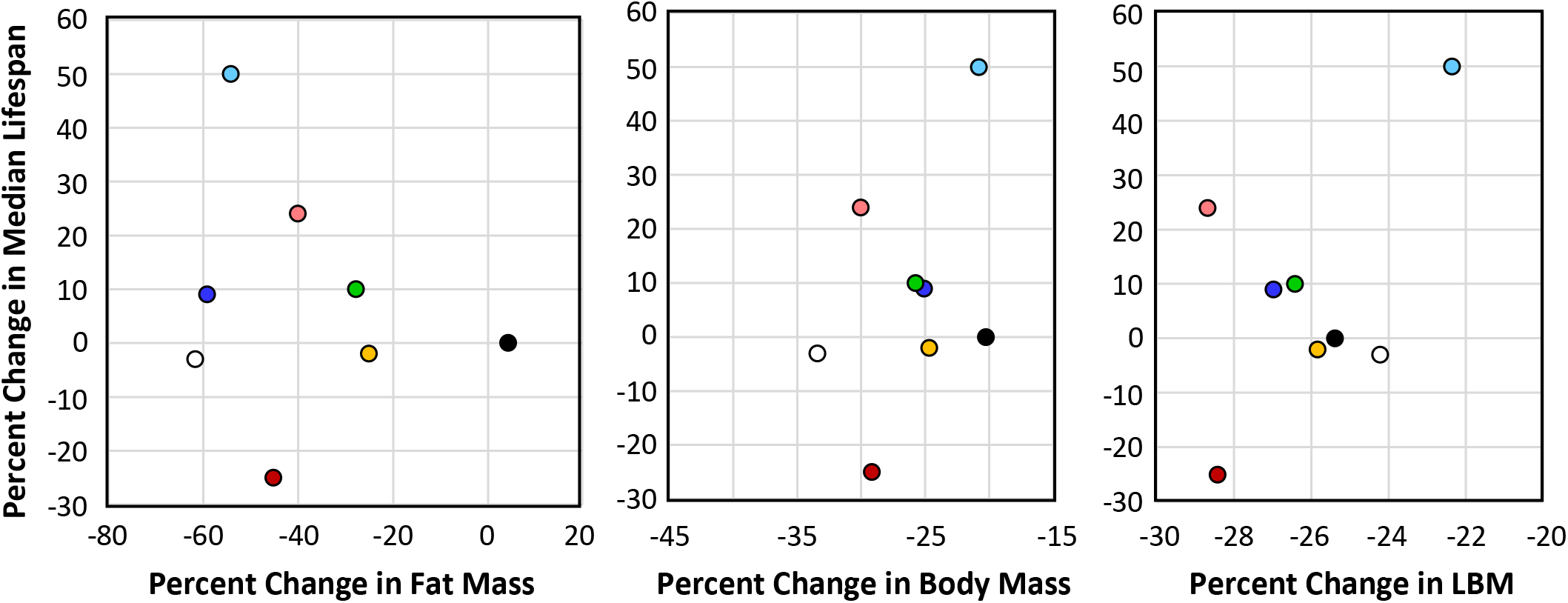
Correlation between changes in body composition and lifespan induced by DR. The average percent change at 12 and 18 months of age in fat mass, body mass, and lean body mass (LBM) induced by 40% DR is plotted versus the change in median lifespan for each of the eight groups of mice: female 115 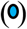, male 115 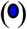, female 107 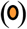, male 107 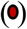, female 98 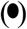, male 98 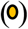, female 97 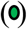, and male 97 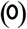.

One of the hallmarks of DR is improved glucose tolerance and insulin sensitivity, and these changes have been proposed to play a role in the life-extending action of DR (Bartke et al., 2001; Barzilai et al., 1998). Therefore, we compared the effect of 40% DR on glucose tolerance in the four RI-lines of male and female mice. Because we have shown that 40% DR can enhance glucose tolerance in C57BL/6 mice within 10 days after implementation of DR (Matyi et al., 2018), we measured glucose tolerance 30 and 90 days after implementing DR (e.g., 2.5 and 4.5 months of age). Figure 4S (in supplement) shows the curves for the glucose tolerance tests and Figure 6 show the data when expressed as the area under the curve. Most of groups showed improved glucose tolerance; however, DR had no significant effect on glucose tolerance in 115-RI females, and in female 97-RI mice glucose tolerance was significantly reduced at 4.5 months of age. Thus there was no relationship between the impact of 40% DR on glucose tolerance and lifespan.

**Figure 6.**
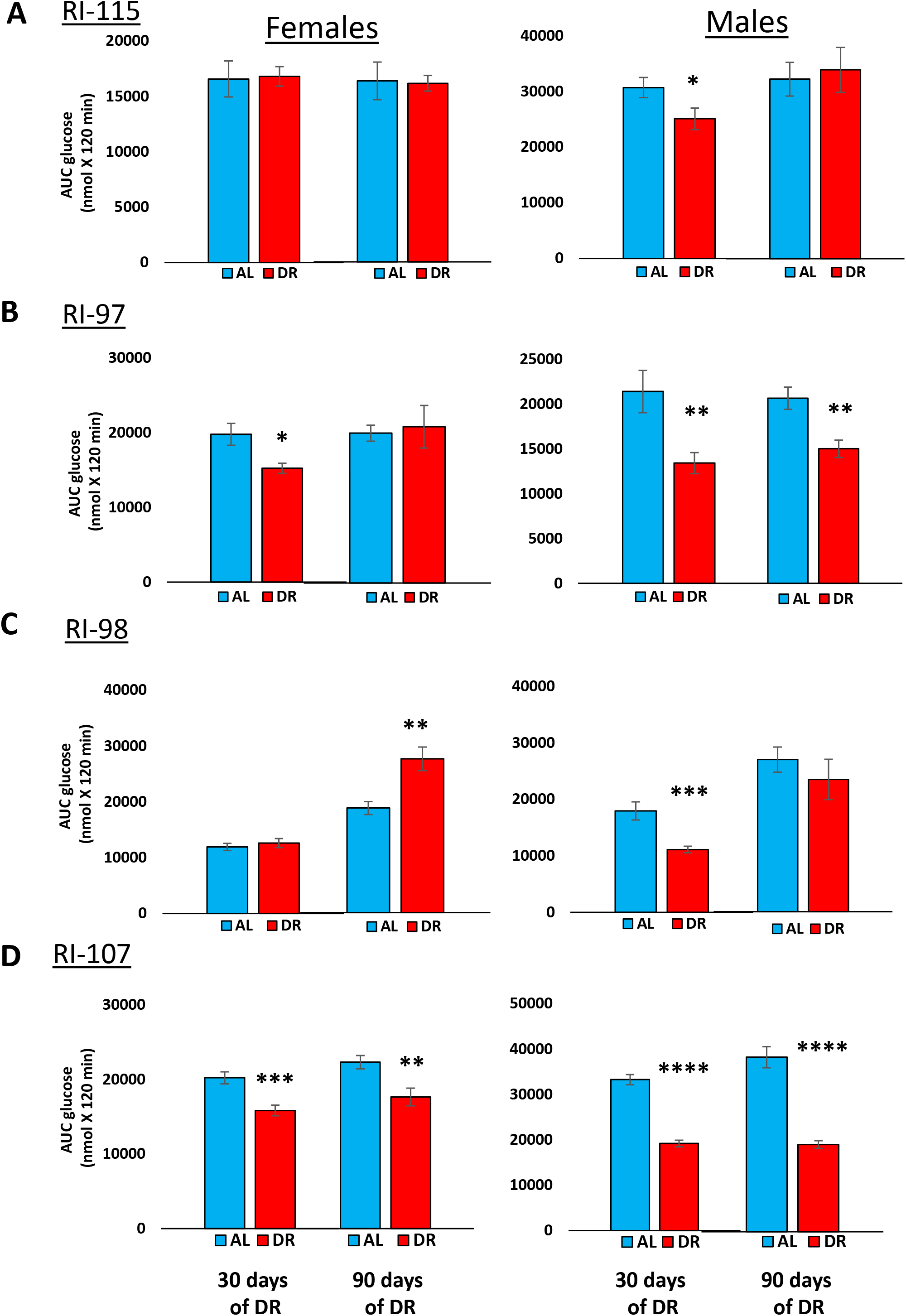
Effect of 40% DR on the glucose tolerance of female and male RI mice. Data from the glucose tolerance curves in Figure 4S in the supplement are expressed as the area under the curve for mice fed AL (blue bars) or DR (red bars) at 3 and 9 months of age. The data are expressed as the mean ±SEM for 9-10 animals per group. **Panel A** 115-RI mice**, Panel B** 97-RI mice, **Panel C**, 98-RI mice, and **Panel D** 107-RI mice. The data for each time point was analyzed as AL Vs DR by two-tailed students t-test. Values where the DR mice (red bars) are significantly different from AL mice (blue bars) are shown by *p>0.05, **p.0.01, and ***p>0.001.

## Discussion

Over the past four decades there have been a large number of studies showing that DR increased the lifespan of different strains of rats and mice. However, there are limited data comparing the effect of DR on different strains of rats or mice under identical conditions conducted by the same laboratory. Turturro et al. (1999), as part of the NIA’s Biomarkers Aging Program, conducted the first study in which the effect of DR on the lifespan was directly compared at the same time in different strains of rats and mice commonly used in aging research and available from the NIA animal colony. These lifespan studies were conducted using an identical degree of DR, 40%. They studied both sexes of three strains of rats (F344, BN, and BNF344F1) and four strains of mice (C57BL/6N, DBA2/N, B6D2F1 and B6C3F1). DR was initiated at 14 weeks of age and was found to increase significantly the lifespans of all strains of female and male rats and mice. For example, the median increase in survival for mice ranged from 15% (male DBA2/N and C57BL/6N mice) to 52% (DBA2/N female mice) with the average increase in median survival of ~30%. Interestingly, the increase in lifespan by DR was similar in males and females for all the rats and the mice, except for DBA2/N mice, which showed a much greater effect of DR in females. In 2010, Liao et al. (2010) conducted a more extensive study of the effect of genotype on the response of mice to 40% DR when initiated at 2 to 5 months of age. They used male and female mice of 41 ILSXISS (formerly called LXS) RI-lines. These inbred RI-mice were generated by Williams et al. (2004) to analyze genetic variation in alcohol sensitivity (Bennett et al., 2006) and were derived from an eight-way cross of the inbred strains: A, AKR, BALB/c, C3H/2, C57BL, DBA/2, Is/Bi, and RIII. As described above, Liao et al. (2010) observed that less than a third of the RI-lines they studied showed a significant increase in lifespan when placed on 40% DR. Of particular interest was the observation that approximately one-third of the mice showed a shortened lifespan on the 40% DR-diet, which was unexpected. The study by Liao et al. (2010) has been the most extensive study to date on the effect of genotype on the life extending action of DR because they used a large number of strains of inbred mice, which were genetically diverse because of the RI-lines came from an eight-way cross. However, because of the large number of RI-lines compared, the study suffered from the small number of mice they used to measure lifespan for each sex and each RI-line.

As noted above, the goal of this study was to determine if the inability of certain RI-lines of mice to respond to DR could be replicated when a larger number (30 to 45 mice/group) of mice were used, which would allow us to detect a 10% change in lifespan (Liang et al., 2003). Of the eight groups of mice (female and male mice of four RI-lines) studied, we were able to replicate the observation of only one of the eight results previously reported by Liao et al. (2010). This was for the male 107-RI mice, which showed that DR resulted in a 22% decreased in mean survival compared to the 50% decrease in mean survival reported by Liao et al. (2010). We did observe that DR significantly reduced the lifespan of male 98-RI mice; however, this decrease (2% for median lifespan) was very small compared to the report by Liao et al. (2021), who observed over a 40% decrease in lifespan. Based on the effect of DR on the lifespan of other strains of mice we studied, which showed similar changes in median survival as the 98-RI mice without significant change in lifespan by DR (e.g., male 97-RI and female 98-RI mice fed 40% DR and male and female 115-RI mice fed 10% DR), we believe that the small effect of DR on the lifespan of male 98-RI mice, even if real, may be less likely to be replicated across a broad variety of environments/laboratories. Because we were unable to replicate the effect of DR in seven out of the eight of the groups of mice that Liao et al. (2010) studied, their data should be viewed cautiously until the lifespan of the other RI-lines are determined using larger numbers of mice.

Although we were unable to replicate the observations reported by Liao et al. (2010) in seven out of the eight groups studied, our data demonstrated that the effect of DR on lifespan varied greatly in the four RI-lines studied. DR significantly increased the lifespan of four of the eight groups, e.g., female and male 115-RI mice and female 97-RI and 107-RI mice. The increase in median lifespan ranged from 10 to 50%. However, DR had little effect (less than 3% change in median survival) on the lifespan of three groups, male 97-RI mice and female and male 98-RI mice. Only the male 107-RI mice showed a major decrease in lifespan. We also observed that DR had quite different effects on lifespan of two of the RI-line. In the 97-RI line, DR increased (10%) the lifespan of female mice but had no significant effect on the lifespan of male mice. In the 107-RI line, the sex difference was major. DR increased the median survival of female mice 24% and reduced the median survival of male mice 22%.

Previous studies have suggested that the ability of mice to preserve their fat mass in response to DR was correlated with a greater increase in lifespan (Liao et al., 2011; Mitchell et al., 2016) and that glucose and insulin sensitivity was important in the anti-aging action of DR (Bartke et al., 2001; Barzilai et al., 1998). Therefore, we measured the changes in body and fat mass and glucose tolerance induced by DR in the eight groups of mice. Table 2 summarizes our findings listing the eight groups of mice in order of the effect of 40% DR on their median survival and the effect of DR on body composition and glucose tolerance. As can be seen from Table 2, we were unable to show any consistent association between the effect of DR on any of these measures and the effect of DR on lifespan.

**Table 2.** Summary of Data from the Four RI-Lines of Mice Studied.

| **RI-Line** | **$\Delta$ in Median Lifespan by DR** | **Median Lifespan of AL** | **$\Delta$ in Body Mass by DR<sup>a</sup>** | **$\Delta$ in Fat Mass by DR<sup>a</sup>** | **$\Delta$ in GTT by DR<sup>b</sup>** |
| --- | --- | --- | --- | --- | --- |
| 115-Female | +50% | 630 days | -21% | -54% | -1% |
| 107-Female | +24% | 846 days | -30% | -40% | -21% |
| 97-Female | +10% | 951 days | -26% | -28% | +4% |
| 115-Male | +9% | 882 days | -25% | -59% | +5% |
| 98-Female | NS | 1062 days | -20% | +4% | +47% |
| 98-Male | NS | 1069 days | -25% | -25% | -13% |
| 97-Male | -2% | 966 days | -33% | -61% | -27% |
| 107-Male | -22% | 930 days | -29% | -45% | -50% |

<sup>a</sup>Average change at 13.5 and 18 months of age. <sup>b</sup>Change measured at 4.5 months of age.

In summary, our study was to a large extend unable to replicate the effect of DR on the lifespan of the four RI-lines reported by Liao et al. (2010); therefore, the lifespan data in their study should be considered suspect because of the limited number of mice used to measure lifespan. However, our data support the general conclusion of their study that genotype has a significant impact on the response of an animal to DR. While we observed half of the groups of mice we studied showed an increase in lifespan when fed DR, the other half either did not respond to DR or showed a decrease in lifespan. These RI-lines are a potentially important resource for investigators studying the anti-aging mechanism of DR because these strains of mice will allow investigators for the first time to compare mice in which DR increases lifespan to mice where DR has either no effect or reduces lifespan. These comparisons will allow investigators to identify pathways that are altered only in mice showing an increase in lifespan, giving us a better understanding of mechanism that is involved in the anti-aging action of DR.

## Experimental procedures

### Animals and Lifespan Analysis

We obtained the following four RI-Lines from The Jackson Laboratory: ILS/ISS115/TejJ (115-IR), ILS/ISS97/TejJ (97-RI), ILS/ISS98/TejJ (98-RI), and ILS/ISS107/TejJ (107-RI) at 4 weeks of age. The mice were housed in the animal facility at the University of Oklahoma Health Sciences Center and maintained under SPF conditions in a HEPA barrier environment. The mice were housed under controlled temperature and light conditions (12-12h light-dark cycle) and fed *ad libitum* irradiated NIH-31 mouse/rat diet from Teklad (Envigo, Madison, WI). At 6 weeks of age, the mice were separated into the different dietary regimens, e.g., *ad libitum* (AL), 10% DR, 20% DR, and 40% DR for the 115 RI-mice and AL and 40% DR for the 97-RI, 98-RI, and 107-RI mice. The food consumption by the AL group of each RI-line and sex was measured every week until 6-months of age and then every month and the amount of NIH-31 diet given to the DR groups each day was adjusted accordingly, i.e., the DR groups were fed 90%, 80% or 60% of the food consumed by the AL mice for 10%, 20%, and 40% DR, respectively. We did not do a step-wise reduction in food given the mice; rather the mice were immediately put on 40% (or 10 and 20%) DR at 6 weeks of age to be consistent with the study by Liao et al. (2010). It should be noted that the DR diets were not fortified with vitamins or minerals, which was identical to the DR protocol used by Liao et al. (2010). The mice in the survival studies were allowed to live out their lifespan without any manipulations except for cage changes every other week and the daily feeding of the DR groups. The mice were housed 5 mice/cage initially and were maintained in their respective cages until they died resulting in less than 5 mice/cage as mice in the cage died. The mice were monitored daily, on weekends, and holidays for overall health and morbidity and allowed to die naturally unless they were either unable to move to obtain food/water, experience pain from the presence of large tumors, or exhibit a major loss of weight indicating they would die within 24 to 48 hours.

The statistical analysis of the lifespan data was conducted by the Comparative Data Analytics Core of the University of Alabama at Birmingham Nathan Shock Center at the Indiana University School of Public Health, and the data can be accessed via the Comparative Data Analytics Core. Using R software, the following analyses were performed to compare the lifespan curves of the AL and DR groups: (1) mixed effects Cox proportional hazards analysis, (2) parametric survival analysis, using exponential, log-normal, Weibull, and Gompertz distributions, (3) Quantile mixed regression to test the difference in median (50^th^ percentile) lifespan, (4) Linear Mixed Models to compare mean lifespans, (5) Kolmogorov-Smirnov tests for the difference in distributions, and (6) the maximum lifespan test (Gao et al., 2008). For the maximum lifespan test the threshold for “long” lifespan was set to be the 90^th^ percentile of all the groups combined, and a new variable Z was coded, where Z=0 for animals dying before the threshold, and Z= lifespan for animals reaching the long-life threshold. Wilcoxon-Mann-Whitney tests then compare Z between groups for significance. Notably, there were no missing (censored) data, which allowed standard comparisons of distributions that do not require accommodation for censored data. The analyses were performed separately for each combination of sex and strain. The AL group was used as a reference group.

To account for potential correlation of animals housed within cage, and animals arriving to the lab within cohort, random effects were included in the analyses for cage and cohort, where cages are nested within cohort. For the mixed effects Cox models and Linear Mixed models, cohort and cage were included as random effects. Quantile mixed regressions, included cage as a random effect, while the additional variance term for cohorts was precluded in standard software (R, SAS, Stata). Parametric survival models, the Kolmogorov-Smirnov test and the Maximum Lifespan test do not include random effects which may inflate type I error. Comparing AICs between the different parametric distributions, the lowest AIC was realized with the Gompertz distribution; therefore, the results with the Gompertz distribution were reported. P-values are all two-sided tests, comparing each diet group to the AL group, and are reported unadjusted.

### Body Composition and Glucose Tolerance Text

Body composition and glucose tolerance was conducted in a separate cohort of mice for longitudinal analysis that were maintained on AL and DR diet for each line and sex. Body composition of the AL and DR fed live mice were measured using nuclear magnetic resonance spectroscopy (NMR-Bruker minispec) at ~2.5, ~4.5, ~13.5, and ~16.5 months of age (30, 90, 360 and 500 days of DR respectively). Body fat and lean body mass of the animals in each group were measured.

Glucose Tolerance was determined on each strain and sex after an overnight fast of mice at ~2.5 and ~4.5 months of age (30 and 90 days of DR respectively). Mice were weighed and injected intraperitoneal with 20% glucose (2g/kg) and blood glucose levels, collected from tail, were measured over a 120-minute period using a glucometer (Contour next EZ from Bayer). The area under curve (AUC) for each curve was determined and represented as AUC glucose (mmol X 120 min).

## Acknowledgements

The research was supported by the following NIH grants: R01AG045693 (AR), KO1AG 056655-01A1 (AU), and the Oklahoma Nathan Shock Center (P30 AG050911) and the UAB Nathan Shock Center (P30 AG050886) and the Nathan Shock Center Coordinating Center (U24 AG056053). Additional support came from the American Federation of Aging Research 17132 (AU), Oklahoma Center for Adult Stem Cell Research (AU) and a Senior Career Research Award 1IK6BX005238 (AR) from the Department of Veterans Affairs.

## Authors’ contributions

AU and AR were involved in the design and implementation of the study, data analysis, interpretation of the results and manuscript writing. SM, KG and MRB were involved in the maintenance of mice colonies and longitudinal body composition analysis. DA, KE, XC and SD conducted the statistical analysis of the lifespan data.

## Data Availability

The statistical analysis of the lifespan data was conducted by the Comparative Data Analytics Core of the University of Alabama at Birmingham Nathan Shock Center at the Indiana University School of Public Health, and the data can be accessed via the Comparative Data Analytics Core.

