## Supporting File for "A Reevaluation of the Effect of Dietary Restriction on Different Recombinant Inbred (RI) Lines of Male and Female Mice"

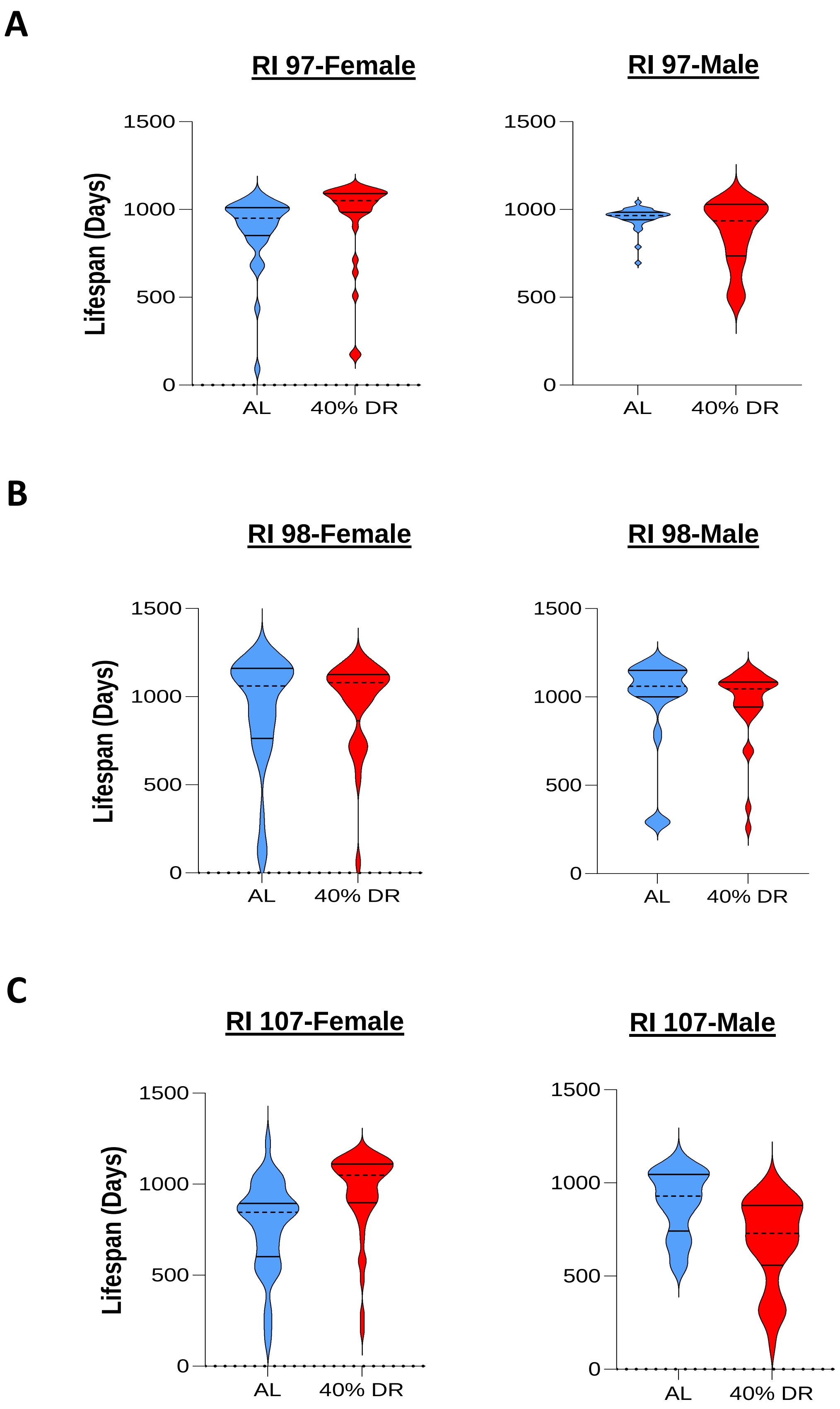


**Figure 1S.** Violin plots for the distribution of the lifespans for each group of mice. The data show the age of death for each mouse in the three RI lines studied: **Panel A**, 97-RI mice, **Panel** B, 98-RI mice, and **Panel C**, 107-RI mice. The solid lines show the quartiles and the dashed line the median.

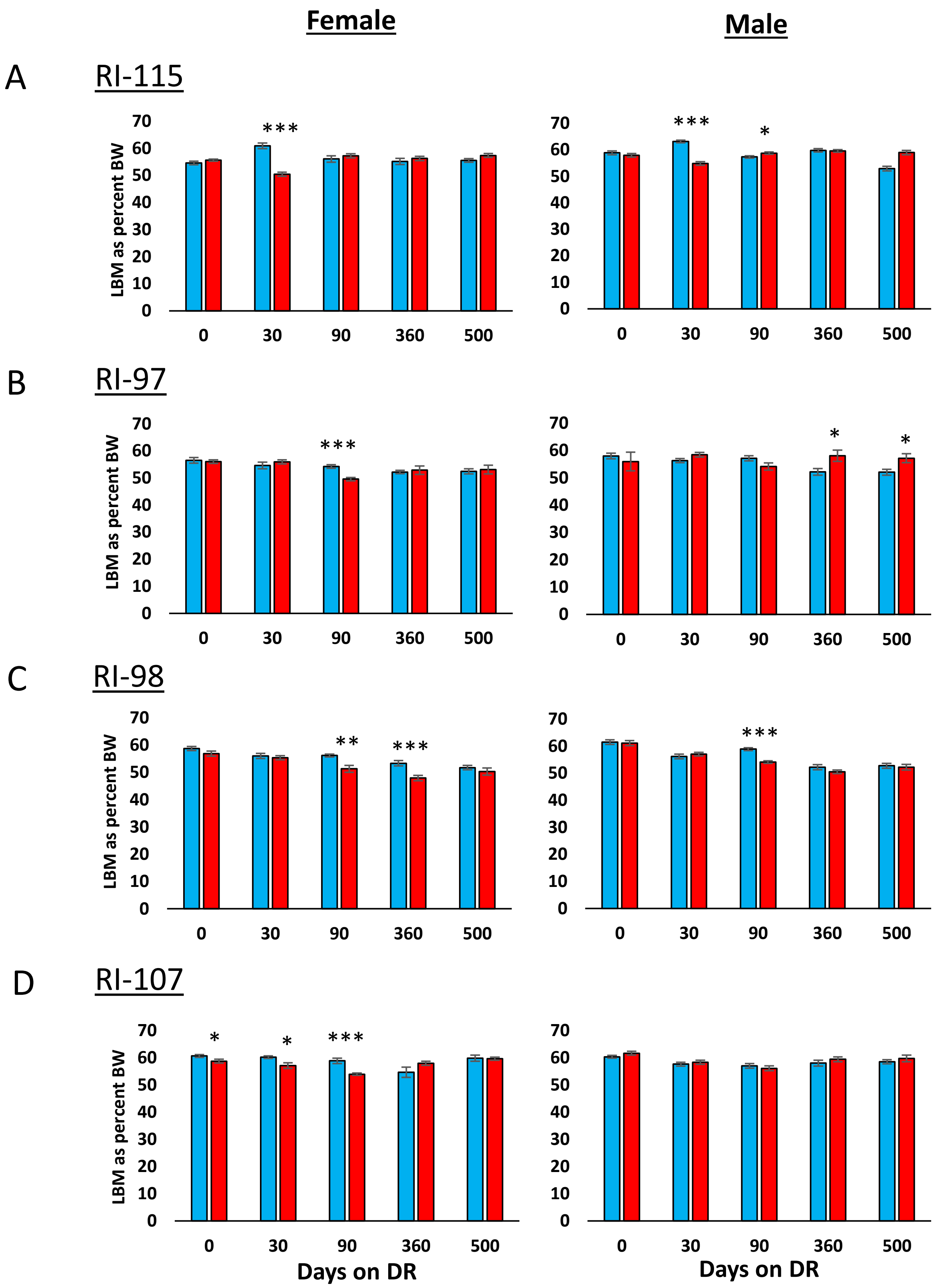


**Figure 2S. Effect of 40% DR on the lean body mass of female and male RI mice.** The percent lean body mass is expressed as the lean body mass divided by the body weight of each animal, and the data show the mean ± SEM for 8-10 animals per group except for 97 Male DR (time points 360 and 500) and 107 male and Female DR (time point 500) groups which have 5 mice. **Panel A** 115-RI mice**, Panel B** 97-RI mice, **Panel C**, 98-RI mice, and **Panel D** 107-RI mice. The data for each time point was analyzed as AL Vs DR by two-tailed students t-test. Values where the DR mice (red bars) are significantly different from AL mice (blue bars) are shown by *p>0.05, **p.0.01, and ***p>0.001.

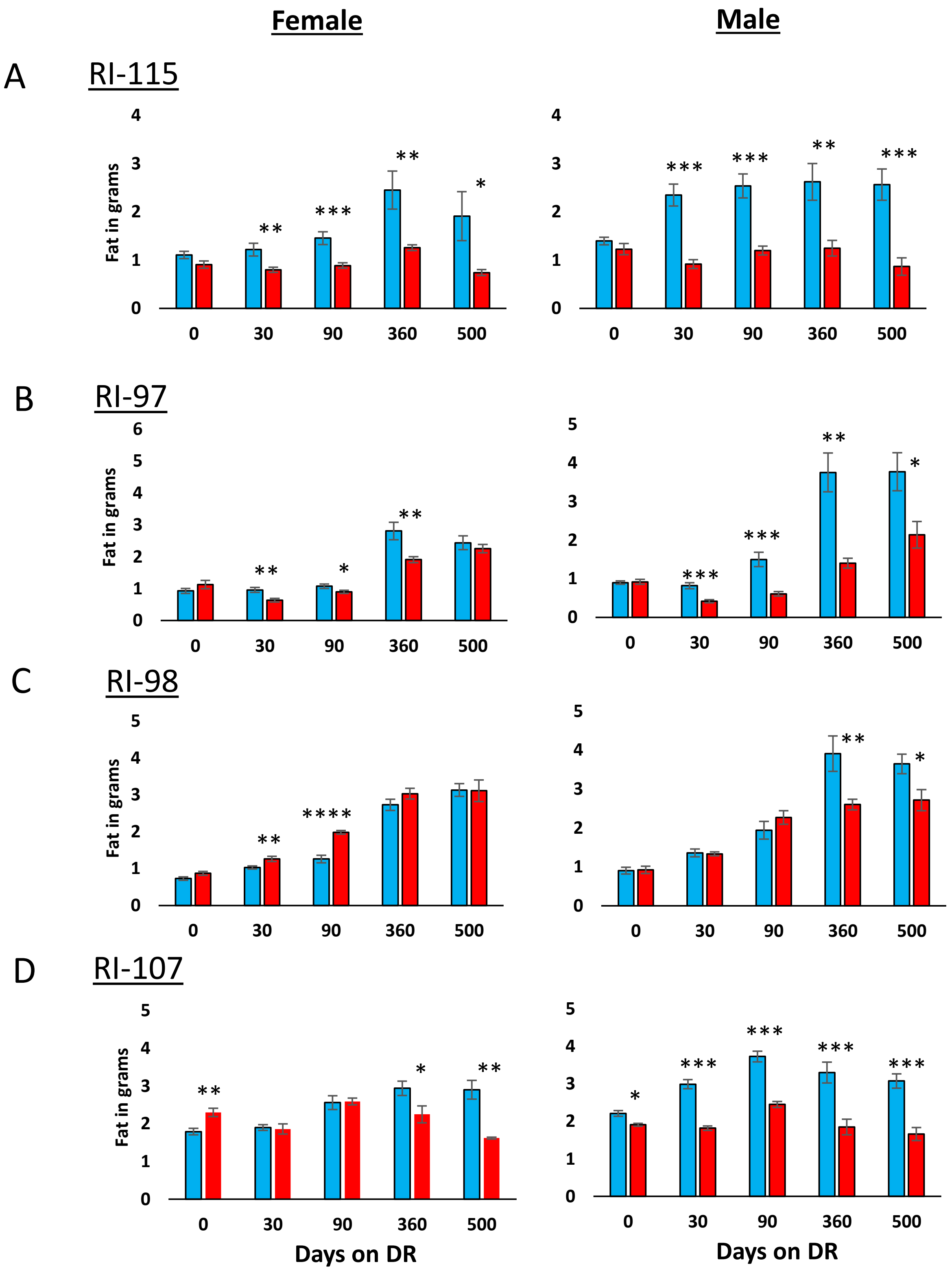


**Figure 3S. Effect of 40% DR on the fat mass of female and male RI mice.** The fat mass is expressed as the mean ± SEM for 8-10 animals per group except for 97 Male DR (time points 360 and 500) and 107 male and Female DR (time point 500) groups which have 5 mice. **Panel A** 115-RI mice**, Panel B** 97-RI mice, **Panel C**, 98-RI mice, and **Panel D** 107-RI mice. The data for each time point was analyzed as AL Vs DR by two-tailed students t-test. Values where the DR mice (red bars) are significantly different from AL mice (blue bars) are shown by *p>0.05, **p.0.01, and ***p>0.001.

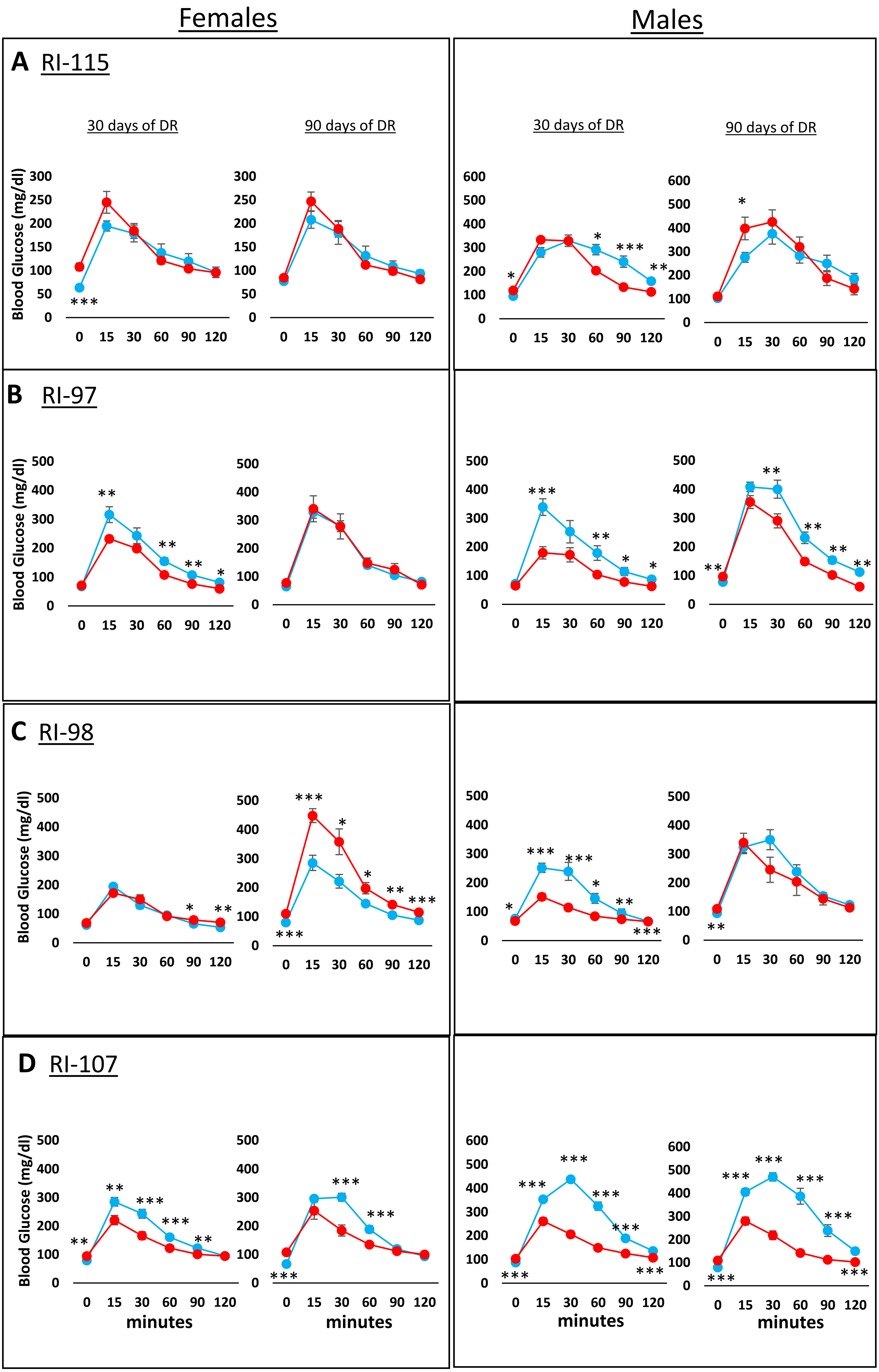


**Figure 4S. Effect of 40% DR on the glucose tolerance of female and male RI mice.** The glucose tolerance curves are shown for 3- and 9-month-old mice fed AL (blue) or DR (red) mice. The data are expressed as the mean ± SEM for 9-10 animals per group. **Panel A** 115-RI mice**, Panel B** 97-RI mice, **Panel C**, 98-RI mice, and **Panel D** 107-RI mice. The data for each time point was analyzed as AL Vs DR by two-tailed students t-test. Values where the DR mice (red curves) are significantly different from AL mice (blue curves) are shown by *p>0.05, **p.0.01, and ***p>0.001.
